# The Secondary Somatosensory Cortex Gates Mechanical and Thermal Sensitivity

**DOI:** 10.1101/2023.05.19.541449

**Authors:** Daniel G. Taub, Qiufen Jiang, Francesca Pietrafesa, Junfeng Su, Caitlin Greene, Michael R. Blanchard, Aakanksha Jain, Mahmoud El-Rifai, Alexis Callen, Katherine Yager, Clara Chung, Zhigang He, Chinfei Chen, Clifford J. Woolf

**Author notes:** Correspondence: Clifford J. Woolf –.

## Abstract

The cerebral cortex is vital for the perception and processing of sensory stimuli. In the somatosensory axis, information is received by two distinct regions, the primary (S1) and secondary (S2) somatosensory cortices. Top-down circuits stemming from S1 can modulate mechanical and cooling but not heat stimuli such that circuit inhibition causes blunted mechanical and cooling perception. Using optogenetics and chemogenetics, we find that in contrast to S1, an inhibition of S2 output increases mechanical and heat, but not cooling sensitivity. Combining 2-photon anatomical reconstruction with chemogenetic inhibition of specific S2 circuits, we discover that S2 projections to the secondary motor cortex (M2) govern mechanical and thermal sensitivity without affecting motor or cognitive function. This suggests that while S2, like S1, encodes specific sensory information, that S2 operates through quite distinct neural substrates to modulate responsiveness to particular somatosensory stimuli and that somatosensory cortical encoding occurs in a largely parallel fashion.

---

Top-down control of somatosensory encoding in the spinal cord by the brain allows for context-dependent modulation of behavioral responses based on changing intrinsic or environmental conditions^1,2^. Somatosensory function relies on this top-down control to accurately predict, evaluate, and appropriately react to mechanical, thermal, and chemical stimuli^2,3^. We and others have found that the primary somatosensory cortex (S1) controls somatosensory reflexive behaviors through excitatory corticospinal neurons that innervate the dorsal horn^4,5^. This constitutes a feed-forward circuit whereby incoming sensory information ascends to S1 for processing and through descending excitatory projections, subsequent information flow in the spinal cord is amplified to facilitate accurate behavioral responses. Inhibition of this S1 corticospinal circuit, therefore, produces decreased mechanical sensitivity^4^. However, S1 does not appear to encode all somatosensory modalities, with a strong bias toward mechanical and cooling inputs and an absence of heat encoding^4,6,7^. This suggests that distinct anatomical regions may participate in the full spectrum of somatosensory top-down control. We sought to identify what other cortical regions that could encode the properties that S1 does not. The secondary somatosensory cortex (S2) is an adjacent cortical region that also processes somatosensory stimuli^8–10^. It has been proposed, based on cortico-cortical and thalamo-cortical neural circuitry, latency of responses, and homology to other sensory cortical areas, that S1 and S2 exist in a hierarchy, with S2 as the higher-order cortical area, processing distinct features of the somatosensory experience similar to the visual system^9–12^. The mouse whisker system has provided some evidence supporting this model and subsets of neurons in S2 appear to encode behavioral choice and recalling of past experience^10,13,14^. However, whether this is the case for somatic stimuli from the body is unclear. S1 can also encode complex aspects of the somatosensory experience questioning a hierarchical relationship and favoring the processing of information in parallel^15,16^. Evidence for parallel information processing, where differential modalities are processed within S1 and S2 is accumulating, particularly in humans^17–19^.

We hypothesized that the secondary somatosensory cortex (S2) may be a crucial substrate for those somatosensory modalities that S1 does not encode. Using a combination of optogenetics and chemogenetics, we now identify the hindpaw area of S2 as a cortical region whose output, in contrast to S1, mitigates mechanical and heat sensitivity. Circuit mapping studies combined with intersectional circuit manipulation strategies identify a higher-order cortical area, the secondary motor cortex (M2), as the target of S2 that governs somatosensory sensitivity. These findings reveal that S2 is a key controller of evoked somatosensory behaviors in a manner quite distinct of that of S1 circuits and dependent on cortico-cortical connectivity to suppress specific somatosensory responses.

## Results

### Inhibition of the Secondary Somatosensory Cortex Enhances Tactile and Thermal Somatosensory Behavioral Responses but Does Not Produce Aversion

To determine if the secondary somatosensory cortex (S2) has a role in somatosensory behaviors we virally targeted expression of channelrhodopsin (ChR2) into parvalbumin (PV) inhibitory interneurons in the hindlimb region of mouse S2 to induce inhibition of the output from S2 (Fig. 1a-c; Supp. Fig. 1)^20,21^. Our viral injection strategy targeted S2 by injecting through the barrel cortex (part of the whisker trigeminal system and independent from the spinal ascending tract) to avoid any bleed through into the laterally adjacent insular cortex which is important for thermosensory perception^6^ (Fig 1a). Implantation of an optical fiber to deliver 40Hz blue light induced strong regional inhibition of the S2 output pyramidal neurons by the activation of PV inhibitory interneurons. Slice recordings from the S2 region confirmed that PV inhibitory interneurons virally transduced with ChR2 are readily excited by blue light and that in consequence, adjacent pyramidal neuron activity is suppressed (see also:^20,21^) (Fig. 1d, e).

**Figure 1:**
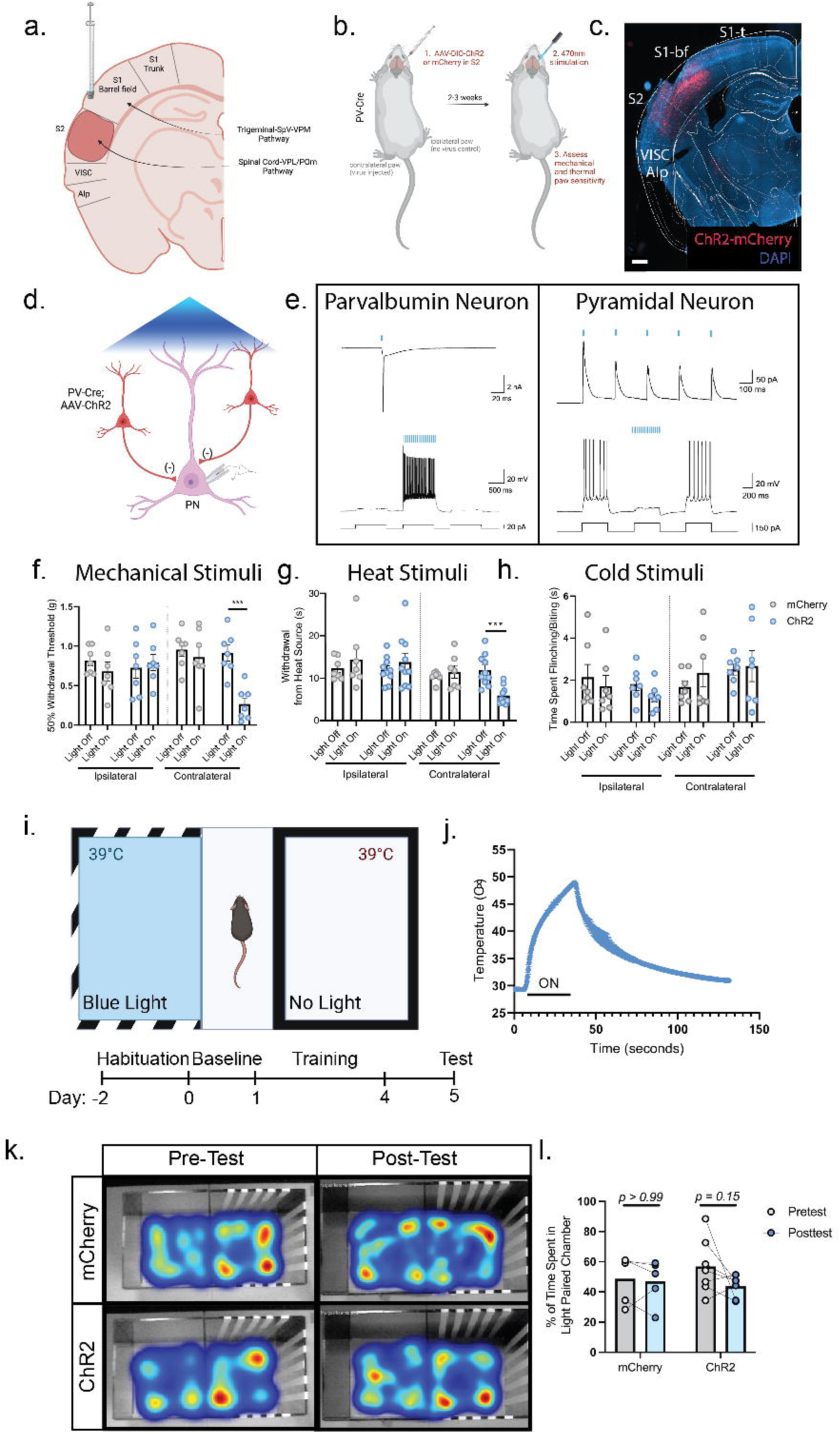
Optogenetic inhibition of the secondary somatosensory cortex (S2) enhances tactile and thermal sensitivity. (a): Schematic diagram of injection strategy into the S2 cortex via needle placement through the barrel cortex. (b): Experimental workflow of optogenetic inhibition of S2. Cre-dependent channelrhodopsin (ChR2) was injected into the S2 region of parvalbumin (PV)-Cre animals. An optical fiber was then placed in the S2 region to enable activation of inhibitory neurons to produce regional net inhibition. (c): Expression of ChR2 virus in PV neurons of the S2 region. Scale bar: 500μm. (d): Schematic diagram depicting slice electrophysiology approach. Pyramidal neurons in S2 were recorded from while adjacent PV-interneurons were activated by blue light illumination. (e): Light-evoked responses in a ChR2^+^ parvalbumin interneuron (left) and a ChR2^-^ pyramidal neuron (right). Left: Blue light stimulation triggers action potential firing in the fast-spiking parvalbumin interneuron in both voltage (up) and current (bottom) clamp modes; Right: Photoactivation of parvalbumin interneurons evokes inhibitory post-synaptic currents (up) and suppresses actional potential firing (bottom) in the pyramidal neuron. (f): Optogenetic inhibition of S2 produces increased mechanical sensitivity in the von Frey mechanical assay. Two Way ANOVA with Tukey’s (n=7 for all groups). (g): Optogenetic inhibition of S2 produces increased sensitivity to heat applied by a Hargreave’s thermal ramp stimulus. Two Way ANOVA with Tukey’s (n=7 mCherry, n=10 ChR2). (h): Optogenetic inhibition of S2 does not affect cold sensitivity in the acetone assay. Two Way ANOVA with Tukey’s (n=7 for both groups) (i): Schematic depicting the conditioned place preference test and experimental timeline. (j): Thermal profile of the heat ramp stimulus applied to the hindpaw. (k): Optogenetic inhibition of S2 does not produce any conditioned aversive behavior. Representative heat maps are color coded to label the most occupied spaces in red and least occupied in blue. (l): Time spent in the paired chamber does not differ between S2 inhibited and non-inhibited animals. Two Way ANOVA with Sidak’s multiple comparisons, p-values as stated (n=5 mCherry, n=7 ChR2). Data presented as mean ± SEM. *p=<0.05, **p=<0.005, ***p=<0.0005.

We then assayed mechanical sensitivity in both the hindpaw contralateral (virus affected) and ipsilateral (unaffected) to the cortical injection site, using graded mechanical von Frey filaments. Stimulation of PV-ChR2 animals with blue light increased mechanical sensitivity compared to PV-mCherry controls (mean: 0.8594g mCherry vs. 0.2659g ChR2) (Fig. 1f). Only the contralateral paw was affected, confirming that there is no effect on the non-injected hemisphere and that this effect is intratelecephalic (mean: 0.6799g mCherry vs. 0.7911g ChR2).

To assess whether the enhanced sensitivity to mechanical stimuli resulting from the inhibition of S2 extends to other somatosensory modalities, we applied a heat ramp until hindpaw withdrawal was observed (Hargreave’s Test). Strikingly, blue light exposure in PV-ChR2 animals but not PV-mCherry control mice elicited a significantly decreased response latency on the contralateral but not the ipsilateral paw, indicating enhanced heat sensitivity (mean: 11.50s mCherry vs. 5.15s ChR2) (Fig. 1g). In contrast, we did not find any behavioral differences in cold sensitivity between PV-ChR2 and PV-mCherry animals (mean: 2.34s mCherry vs. 2.67s ChR2) (Fig. 1h). Therefore, inhibition of hindlimb S2 by activation of PV inhibitory interneurons produces an increase in tactile and heat sensitivity, suggesting that outputs from S2 gate these sensory responses in a manner distinct from S1 which governs mechanical and cold sensory responses.

Based on the neural connectivity of rodent and primate S2 and activity recordings in S2, it might be expected that this cortical region is actively involved in the processing of the aversive components of somatosensory behavior^22–25^. We tested whether S2 output inhibition by PV inhibitory interneuronal activation results in a change in aversive behavior, using a conditioned place aversion assay (Fig. 1i). Mice were conditioned to associate optogenetic inhibition of S2 output with a certain colored chamber (striped vs. solid walls). The floors of both chambers were set at 39°C, the measured average withdrawal-evoking temperature in ChR2-stimulated mice (Fig. 1j). However, after associative training, we observed no aversive response to S2 output inhibition, with mice preferring both chambers equally (Fig. 1k, l) (% time spent in conditioned chamber: mCherry pre-conditioned 48.69% vs. post-conditioned 46.84% and ChR2 pre conditioned 56.90% vs. post-conditioned 43.69%). This suggests that while inhibition of S2 can increase sensitivity to mechanical and thermal evoked behaviors it is not driving an aversive component.

To rule out that S2 is induces an anxiolytic-like behavior which alters somatosensory reactivity, we used an elevated plus maze assay, in which the preference of mice to open or closed arms of an elevated platform are evaluated. This revealed that S2 inhibition did not induce any change in preference to the closed arms compared to mCherry controls (% time spent in open arms mCherry 10.30% vs 4.38% ChR2) (Supp. Fig. 2). This suggests that S2 is primarily involved in the modulation of behavioral responses to somatosensory inputs but not in an affective or aversive fashion.

**Figure 2:**
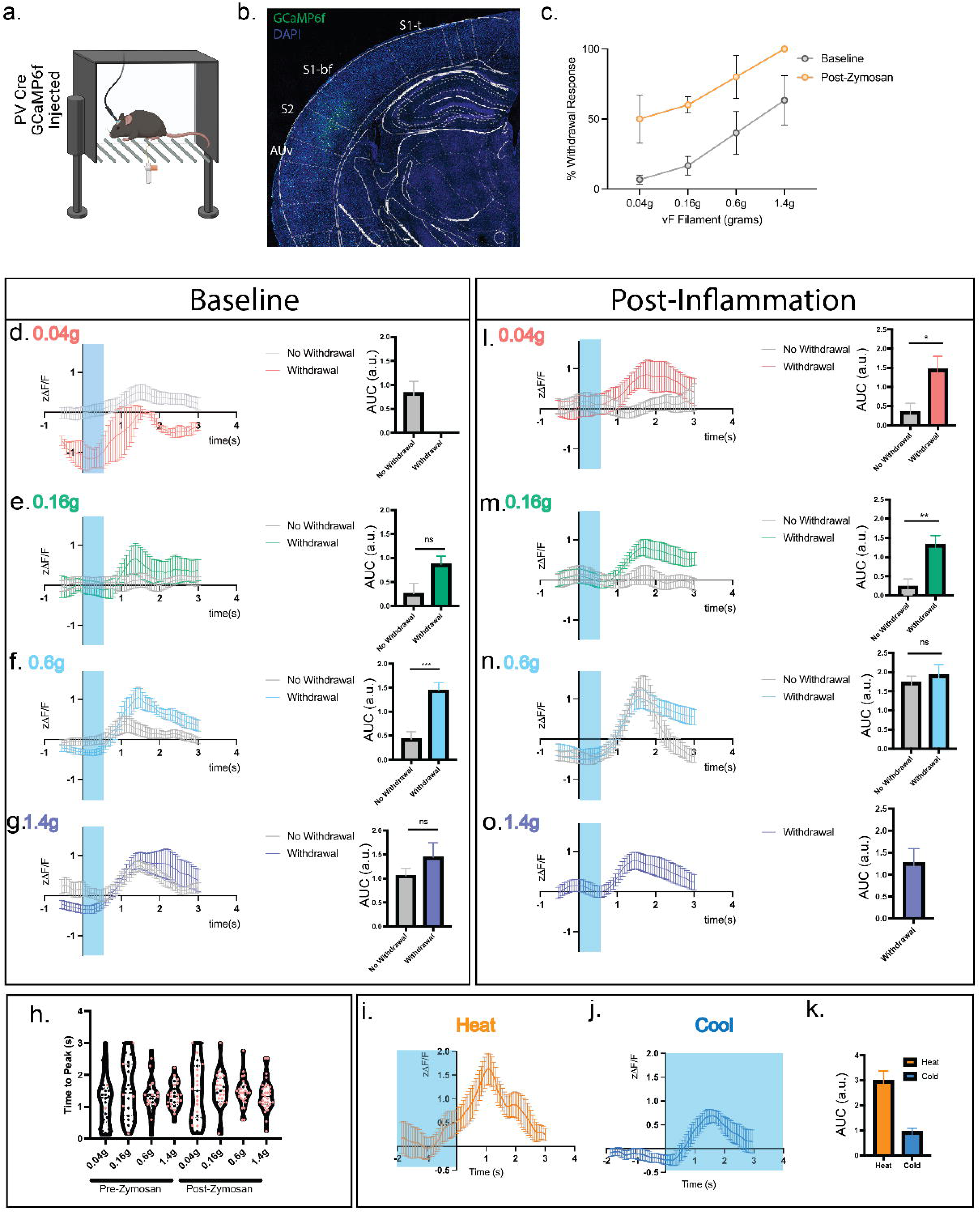
The S2 cortical responses to mechanical and thermal somatosensory stimuli in the non-noxious and noxious range correlates with behavioral outputs. (a): Diagram depicting the experimental strategy of *in vivo* calcium imaging with fiber photometry. GCaMP6f was injected into the S2 region of parvalbumin-cre (PV-Cre) mice, and a fiber lowered into S2 to capture calcium transients. (b): Example of GCaMP6f expression in PV neurons of the S2 region. (c): Percent withdrawal to differentially weighted mechanical von Frey stimulation of the hindpaw in fiber implanted animals both before and after inflammatory induction. (d,e,f,g): Left: Calcium responses of PV neurons in S2 to a 0.04, 0.16, 0.6 and 1.4g mechanical stimulus plotted by Z-scored delta F/F following a single stimulation of the hindpaw at time 0. Blue shading represents the average temporal extent of stimulation. Right: Area under curve analysis of calcium transients produced in d, e, f, g. Unpaired t-test. (h): Time to peak of calcium response to a single mechanical stimulus of the hindpaw. Black circles and red circles represent no paw withdrawal and paw withdrawal trials, respectively. Data shown as a violin plot with the median and interquartile range depicted. (i): Calcium responses in PV neurons in S2 plotted by Z-scored delta F/F to a heat ramp stimulus. Time 0 is time of paw withdrawal from the heat source. Blue shading represents when the heat stimulus is on (j): Calcium responses in PV neurons in S2, plotted by Z-scored delta F/F, to application of acetone to the hindpaw at time 0. Blue shading represents the temporal extent of the acetone stimulus (k): Area under curve analysis of the calcium transients produced in i and j. (l, m, n, o): Left: Calcium responses of PV neurons in S2 plotted by Z-scored delta F/F following a single mechanical stimulation of the hindpaw at time 0, 4 hours post zymosan injection. Force of stimulation in the graphs. Right: Area under curve analysis of calcium transients produced in l, m, n, o. Unpaired t-test. For all experiments, n=3 animals, 5-10 sensory trials per mouse, see Methods. Data presented as mean ± SEM. *p=<0.05, **p=<0.005, ***p=<0.0005.

### Parvalbumin Neurons in S2 are Tuned to Specific Intensities and Modalities of Sensory Input

As inhibition of S2 pyramidal neuron output by activation of parvalbumin (PV) inhibitory interneurons increases somatosensory thresholds, we hypothesized that they may function as a gate that governs the behavioral responses to mechanical and thermal somatosensory stimuli. To test this, we used *in vivo* calcium imaging in awake mice by virally targeting the S2 PV inhibitory interneurons with the calcium indicator GCaMP6f and performing fiber photometry (Fig. 2a, b). Implanted mice had normal somatosensory mechanical detection thresholds, with about 40% of the mice responding to stimulation with a 0.6g and 60% to a 1.4g mechanical force (Fig. 2c). Analyzing those trials in which mice displayed a behavioral withdrawal response revealed that in these cases, S2 PV neurons respond with robust transients 1.3-1.5 seconds after 0.16g through 1.4g mechanical force application with time 0 reflecting the first contact of the filament with the paw (Fig. 2d-g). These temporal dynamics are in line with other fiber recordings made during paw stimulation using GCaMP6f ^26–29^. In the trials in which the mouse did not withdrawal its paw within the 0.04-0.6g range, had in contrast, no calcium transient response, indicating that PV activity correlates with the behavioral response (Fig. 2d-g). However, once in the noxious range of 1.4g, calcium responses in the S2 PV neurons occurred even in the absence of a paw withdrawal, indicating that S2 activity and behavioral response are not always coupled once high-intensity stimuli are detected (Fig. 2g). This suggests top-down circuitry from S2 is able to suppress reflexive behavioral responses to a certain degree of stimulus intensity (largely gating low threshold mechanical stimuli) before spinal reflexes become dominant. Interestingly, the variance in the peak time of individual trials decreased with higher force stimulation (Fig. 2h), with 1.4g producing the most temporally coherent response. This suggests that PV interneurons are more tuned to specific mechanical forces on the paw and fire more synchronously at these forces.

Examination of the calcium transients elicited by PV interneurons during a heat ramp stimulus applied until hindpaw withdrawal, also showed a significant increase in activity beginning ∼0.72 seconds prior to the paw withdrawal event and peaked ∼1.08 seconds post paw withdrawal (Fig. 2i, l). The amplitude of the response to heat was consistently higher than that for mechanical stimuli suggesting a preference of PV+ neural recruitment in S2 to noxious heat stimuli (Fig. 2i, l). Further, the calcium transients in S2 PV neurons increased almost a full second prior to the motor withdrawal response being initiated, indicating that the activity recorded in these inhibitory interneurons is due to sensory input rather than a response to motor output (Fig. 2i).

We also examined the responsivity of S2 to cold stimuli by applying the evaporative coolant acetone and found that S2 is responsive to cold with similar dynamics to that of a 0.6g mechanical stimuli, both in amplitude and timing, with transients peaking ∼1.5 seconds following the stimulus application but are much less responsive than to a heat stimulus (Fig. 2j). These results indicate that S2 PV interneurons are responsive to a broad set of mechanical and thermal somatosensory stimuli and are particularly tuned to low intensity mechanical and heat stimuli. Together with the results that increasing PV activity produces mechanical and heat behavioral hypersensitivity, suggests that S2 acts as a controller of behavioral reactivity to somatosensory stimuli.

Based on this optogenetic and fiber photometry data, we reasoned that increasing peripheral inputs in the setting of peripheral inflammation could shift the tuning of S2 PV neurons and modify their gating properties. To assess this, we performed fiber photometry during the peak phase of an inflammatory insult to the hindpaw in which the fungal ligand zymosan was injected ^30^. Inflammation induced by zymosan significantly shifted the mechanical sensitivity of PV interneurons to a more hypersensitive state (Fig 2l-o) with the neurons responding to lower mechanical forces than at baseline. In contrast to the basal state, low intensity 0.04g stimuli now elicited a calcium response in the S2 PV neurons and the 0.16g response was enhanced (Fig. 2l). Further, the threshold at which behavioral and S2 calcium responses become uncoupled (in which we hypothesize spinal reflexes become dominant) also shifted, and 0.6g stimuli now elicited a response in S2 inhibitory interneurons regardless of the behavioral response (Fig. 2n). Additionally, the time to rise of the transients became more temporally coherent for the 0.16g stimulus, suggesting that PV neurons now alter their tuning properties to now respond to lower threshold stimuli (Fig. 2h). In response to a heat ramp stimulus, the calcium trace from the S2 PV neurons was shifted temporally and its magnitude reduced, although not significantly (Supp. Fig 3). However, this could be due to a change in temporal summation of the heat signal as mice maintain exposure of their paw to the stimulus for only a few seconds before withdrawing.

**Figure 3:**
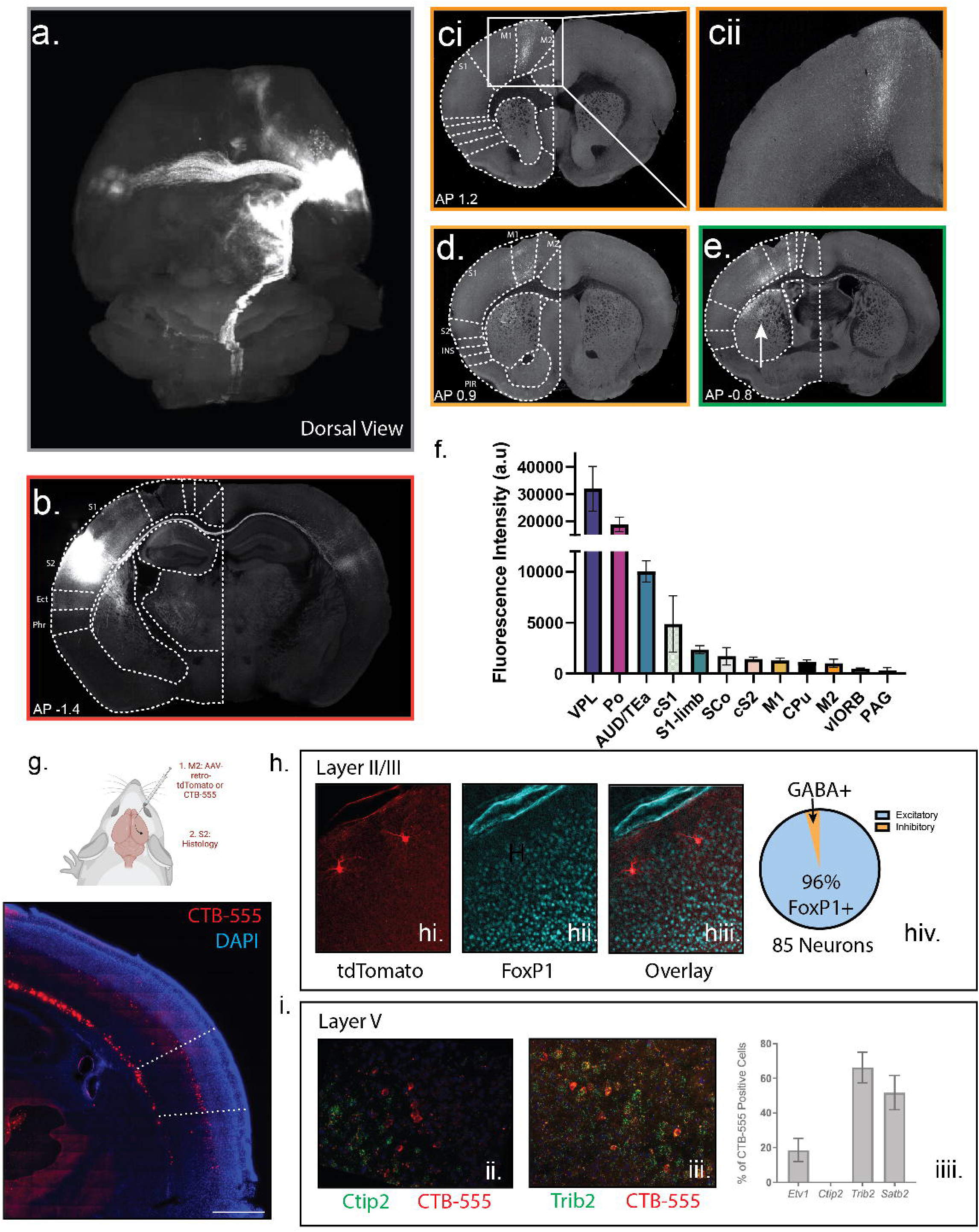
The S2 projectome reveals specific cortical and subcortical sensorimotor targets including the secondary motor cortex (M2). (a): Dorsal view of the reconstruction of S2 projection sites. Tract from S2 to M2 highlighted by an orange arrow. (b): Example S2 injection site. (c): (i): M2 projections (ii): magnified M2 projections. (d): Projections to the M1 cortex. (e): Projections to the primary somatosensory cortex (S1). Arrow denotes projections to the striatum. (f): Average intensity of projections to different brain regions. S2 – red (contralateral light red), S1 – green (contralateral light green), vlORB – navy, M1 – yellow, M2 – orange, AUD/TEa – teal, caudate putamen – black, Po of thalamus – purple, VPL of thalamus – blue, superior colliculus – white, PAG – dark purple. (g): Top: Schematic for ascertaining the identity of S2-to-M2 neurons by injecting either the retrograde dye CTB-555 or AAV2/retro-CAG-tdTomato into M2 and analyzing the S2 region. Bottom: Representative image of the S2 region labeled with CTB-555. Scale bar: 500μm (h): Identity of Layer II/III neurons. (hi): Image of retro-CAG-tdTomato labeled neurons in layer II of S2. (hii): Foxp1 staining in the cortex as a marker of excitatory neurons. (hiii): Merged image reveals overlap between retro-CAG-tdTomato labeled neurons and Foxp1. Analysis of 3 animals with 85 neurons total. (i): Identity of Layer V neurons by RNAscope. (ii): Example of non-overlap between CTB-555 (red) labelled neurons and Ctip2 RNAscope probe (green). (iii): Example of overlap between CTB-555 (red) labelled neurons and Trib2 (green). (iiii): Quantification of overlap between CTB-555 labelled neurons and Ctip2, Etv1, Satb2, and Trib2. n=3 animals. Data presented as mean ± SEM.

### Anterograde Viral Tracing from S2 Reveals Cortico-cortical and Subcortical Innervation with Distinct Innervation of Secondary Motor Cortex (M2)

Unlike the primary somatosensory cortex (S1), S2 does not project to the dorsal horn of the lumbar region of the spinal cord where hindpaw sensation and movement is generated^5,31^. We hypothesized that a novel circuit must be responsible for altering the stimulus-evoked somatosensory behavior. To define this circuit, we injected an AAV encoding tdTomato under the CAG promoter across all layers of the S2 cortex and performed serial 2-photon tomography^32^. We then aligned these images with the Allen Brain Atlas common framework^33^ to identify local and long-distance projection targets (Fig. 3a – quantified in Fig. 3f). We found that S2 makes significant cortical and subcortical projections, the densest of which were observed subcortically in the ventrolateral (VPL) and posterior complexes (Po) of the thalamus, where limb sensory information is received from the spinal cord^34^. These are likely both feedback projections and part of the cortico-thalamo-cortical loop circuitry^12,34^. Other subcortical projection targets included the superior colliculus (SCo), the caudate putamen (CPu) (shown in Fig. 3c, d, and e (arrow)), the periaqueductal gray (PAG), and cervical corticospinal tract, all of which have been implicated in somatomotor circuitry^35^. S2 also made significant corticocortical projections, the densest of which were local to the adjacent auditory/temporal association cortices (AUD/TEa). As observed by others, S2 made significant connections to the contralateral S2, as well as ipsilateral/contralateral projections to the primary somatosensory cortex (S1) (Fig. 3a, e) and to the ipsilateral primary motor cortex (M1)^36^ (Fig. 3d). We also observed a major projection to the ipsilateral prefrontal cortex, specifically the secondary motor cortex (M2)^37^ (Fig. 3a (orange arrow) Fig. 3c and cii). These projection targets place S2 within the common somatomotor architecture^38^, but also reveal engagement with diverse neural substrates capable of transducing and calculating sensory information.

The secondary motor cortex (M2) receives sensory inputs and exerts complex effects on behavior. Indeed, during a sensory decision-making task, widefield imaging demonstrates sensory to frontal waves of activity that occurs before a behavioral choice^39–41^. Further, lesioning or silencing of M2 produces alterations in this behavioral choice^42–45^. We therefore hypothesized that S2 connectivity with the M2 region might underlie the observed somatosensory sensitivity where behavioral choice is altered to a hypersensitive state.

We first anatomically defined the neural identity of S2 to M2 projection neurons by injecting either a retrograde dye (CTB-555) or sparse-labeling with retrograde AAV-tdTomato into M2 and performed RNAscope *in situ* hybridization and immunofluorescence using defined markers (Fig. 3g, top). The majority of S2-to-M2 neurons were located in layer V, specifically Va, with scattered neurons throughout layer II/III (Fig. 3g, bottom). Using four markers of layer V neurons, we found that S2-to-M2 neurons are positive for the callosal intratelencephalic (IT) excitatory neuron markers *Trib2*, *Etv1*, and *Satb2*, but negative for *Ctip2*, a marker of layer Vb excitatory corticospinal neurons (Fig. 3h)^46–48^. Further, the scattered neurons within layer II/III were also found to be excitatory as they express FoxP1, a marker of cortical excitatory neurons (Fig. 3i)^49^. This agrees with tracing data from the Allen Brain Atlas in which anterograde tracing of excitatory layer V neurons in S2 contributes to the largest population of M2 projections with some projections from layer II/III^50^ (Supp. Fig. 4). S2 to M2 projecting neurons are, therefore, excitatory intratelecephalic (IT) projection neurons, largely originating from layer Va.

**Figure 4:**
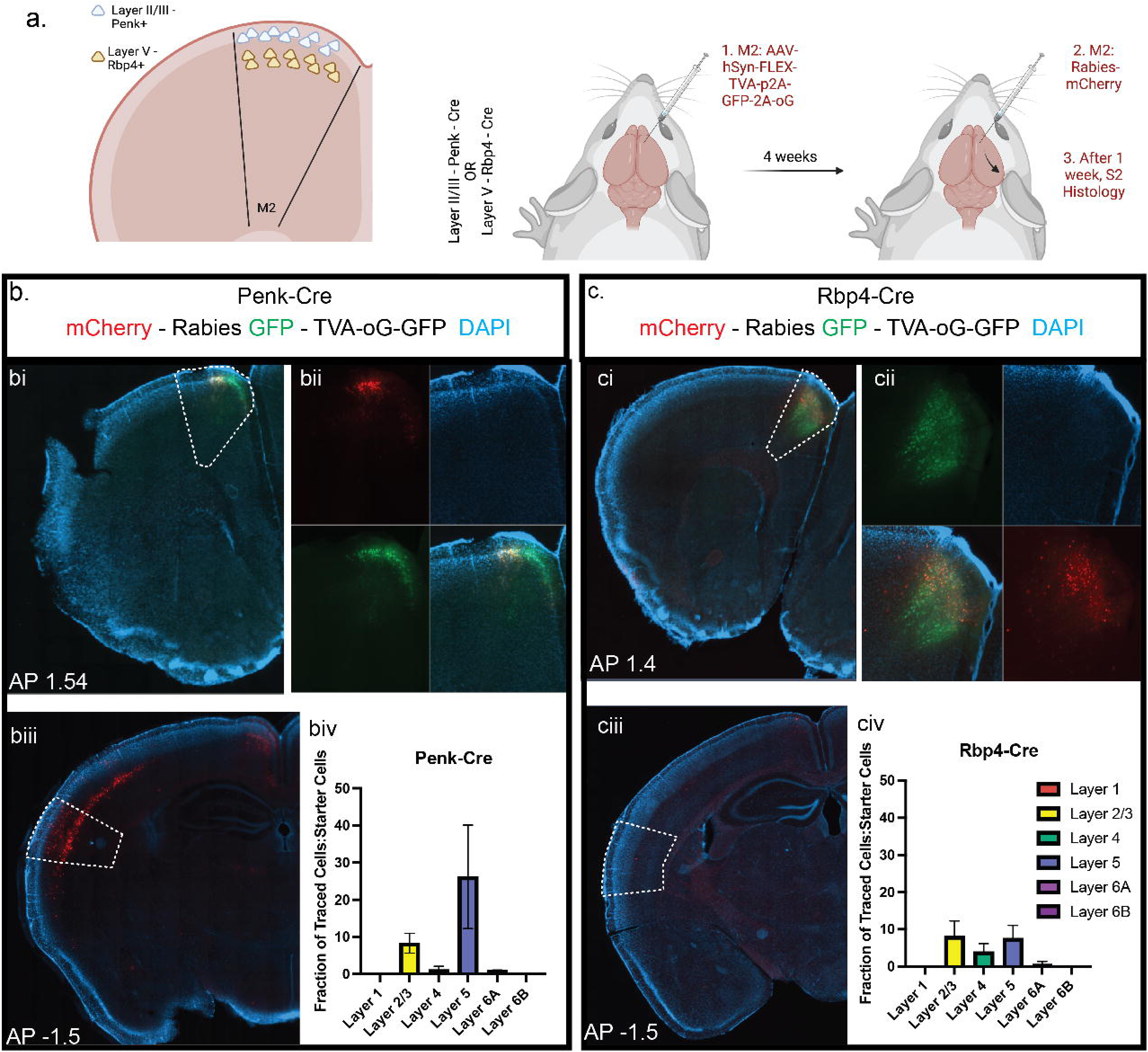
S2 monosynaptically connects with upper layer II/III neurons in M2. a. Schematic diagram of layer specific rabies tracing. b. Example of M2 injection site in Penk-Cre animal. GFP: AAV-hSyn-Flex-TVA-p2A-GFP-2A-oG. mCherry: Rabies. Blue: DAPI. bi. Widefield view of M2 with M2 region outlined. bii. Zoomed in view divided by channel. biii. Widefield view of S2 with S2 region outlined. biv. Layer-specific quantification of the percentage of “starter cells” in M2 compared with “input neurons” in S2 in Penk-Cre Mice. c. Example of M2 injection site in Rbp4-Cre animal. GFP: AAV-hSyn-Flex-TVA-p2A-GFP-2A-oG. mCherry: Rabies. Blue: DAPI. ci. Widefield view of M2 with M2 region outlined. cii. Zoomed in view divided by channel. ciii. Widefield view of S2 with S2 region outlined. civ. Layer-specific quantification of the percentage of starter cells in M2 compared with traced neurons in S2 in Rbp4-Cre Mice. Data presented as mean ± SEM. For all experiments, n=3 mice.

### Layer Specific Connectivity Reveals Hierarchical Relationship Between S2 and M2

The layered structure of the cortex is organized in a manner to facilitate differential computations. Cortico-cortical communication pathways can stem from layer II/III to layer II/III or from deeper layer IT neurons targeting superficial layers^38,51,52^. Modulatory pathways typically innervate the superficial layers whereas driver pathways tend to innervate deeper layers^52,53^ (but see^54^). We set out to address which layer of M2 is innervated by S2 with monosynaptic rabies tracing using Cre drivers that express in layer II/III (Penk-Cre) or layer V (Rbp4-Cre) (Fig. 4a). In this strategy, Cre-positive neurons act as “starter cells” that are selective hosts of rabies infection and retrograde monosynaptic transport. This identifies “input cells”, thereby providing direct evidence of monosynaptic connectivity between two neurons (Fig. 4a). Viral injections into M2 successfully targeted layer II/III and layer V neurons, respectively, labelling many “starter cells” (Fig. 4b (bi-bii), c (ci-cii)). Examination of the rabies virus mCherry+-labelled “input cells” in S2 compared to the total “starter cell” number in M2 revealed the majority of the S2-to-M2 projection neurons in reside in deeper layers of S2 (notably layer V in line with our previous retrograde tracing) and that they primarily innervate superficial layer II/III Penk+ neurons in M2 (Fig. 4 biii-biv and ciii-civ). This arrangement is in line with the role of these neurons playing a modulatory role on M2 activity. Recent M2 single cell sequencing studies have identified layer II/III Penk+ neurons as excitatory neurons ^55,56^. This suggests that inhibition of S2 during somatosensory stimulation, results in decreased M2 activity, and compatible with the recent findings that the net effect of S1/S2 inhibition on L2/3 neurons in M2 is inhibitory ^57^.

### Chemogenetic Inhibition of S2-to-M2 Projecting Neurons Enhances Tactile and Thermal Sensitivity

To functionally manipulate S2-to-M2 IT neurons we used an intersecting viral strategy with injection of a retrograde AAV encoding Cre-recombinase into the M2 region along with an AAV encoding either a Cre-dependent excitatory (HM3Dq) or an inhibitory (hM4Di) chemogenetic receptor (designer receptor activated exclusively by designer drugs (DREADDs)) to either activate or inhibit S2-to-M2 projection neurons with the ligand clozapine n-oxide (CNO) ^58,59^ (Fig. 5a, b). CNO application rapidly and reversibly inhibited firing in S2-to-M2 projection neurons in slices from the inhibitory DREADD mice (Fig. 5c). Likewise, analysis of c-Fos expression as a marker of activity-dependent early immediate gene transcription demonstrated that CNO injection significantly upregulated c-Fos expression in animals injected with a Cre-dependent excitatory DREADD (HM3q) but not a mCherry virus (Supp. Fig. 5a-d). This demonstrates our chemogenetic approach can increase or decrease S2-to-M2 neural activity efficiently.

**Figure 5:**
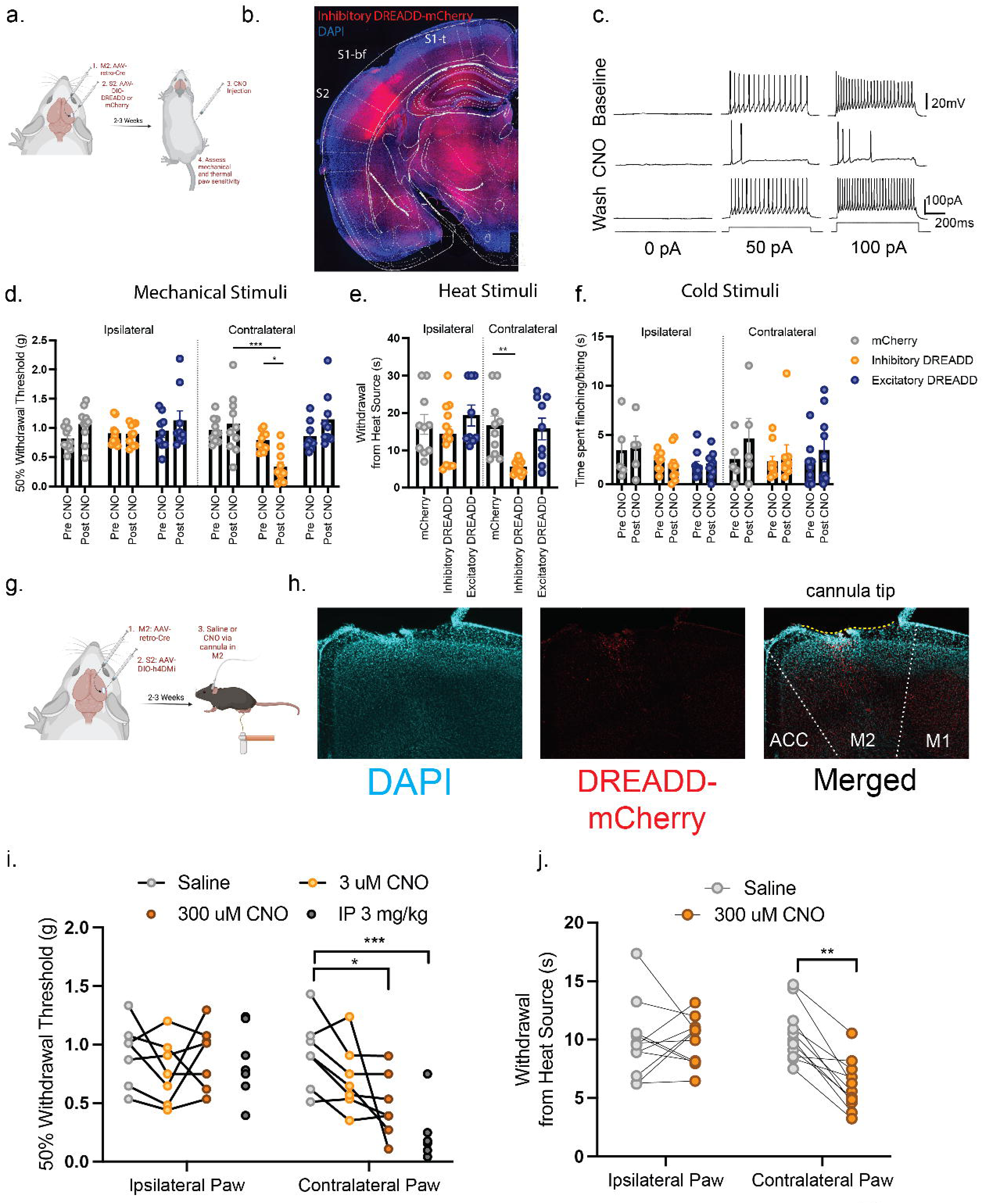
Chemogenetic inhibition of S2-to-M2 projecting neurons enhances mechanical and heat sensitivity. (a): Schematic diagram and timeline of chemogenetic manipulation of S2-to-M2 projecting neurons. (b): Representative images of S2 region following injection AAV-DIO-Inhibitory DREADD in S2 and AAV2/retro-Cre in M2. (c): CNO ligand administration in *ex vivo* brain slices during patch clamp recording inhibits neurons infected with inhibitory DREADDs. (d): Chemogenetic inhibition of S2-to-M2 neurons produces tactile sensitivity in the von Frey assay. mCherry: n=10, Inhibitory DREADD: n=11, Excitatory DREADD: n=10. Two-Way ANOVA followed by Tukey’s multiple comparisons test mCherry contralateral vs. Inhibitory DREADD contralateral *p=0.0002*. (e): Chemogenetic inhibition of S2-to-M2 neurons produces heat sensitivity in the Hargreave’s assay mCherry: n=10, Inhibitory DREADD: n=11, Excitatory DREADD: n=10. Kruskal-Wallis H=25.08 *p=0.0001* Dunn’s multiple comparisons mCherry contralateral vs. Inhibitory DREADD contralateral *p = 0.0032* (f): Chemogenetic inhibition or excitation of S2-to-M2 neurons produces no effect on acetone-induced cold sensitivity mCherry: n=5, Inhibitory DREADD: n=6, Excitatory DREADD: n=10 Two-Way ANOVA followed by Tukey’s post-hoc test. (g): Schematic diagram of chemogenetic inhibition of M2 projections via cannula administration of CNO. (h): Representative image of cannula placement above M2. (i): Chemogenetic inhibition of local M2 projections reproduces the tactile sensitivity observed with systemic CNO application. mCherry: n=8, inhibitory DREADD: n=8 Contralateral Paw Saline vs. 300µM CNO Two Way ANOVA followed by Tukey’s. (j): Chemogenetic inhibition of local M2 projections reproduces the heat sensitivity observed with systemic CNO application. mCherry: n=8, inhibitory DREADD: n=8. Contralateral Paw Saline vs. 300µM CNO Two Way ANOVA followed by Sidak’s. Data presented as mean ± SEM. *p=<0.05, **p=<0.005, ***p=<0.0005.

Examining mechanical and thermal sensitivity in these animals revealed that injection of animals that express the inhibitory DREADD in the S2-to-M2 projection neurons with CNO phenocopied our behavioral observations with optogenetic inhibition of S2 by activation of PV+ inhibitory interneurons. Specifically, 30 minutes after CNO injection, S2-to-M2 inhibitory DREADD animals showed strong tactile hypersensitivity (hM4Di post-CNO contralateral (0.5534g) paw vs. ipsilateral (0.8936g)) and heat sensitivity (hM4Di contralateral paw 6.007s vs. mCherry contralateral paw 16.53s) (Fig. 5d, e). Again, cold sensitivity remained unaffected (mCherry contralateral 2.480s vs. hM4Di contralateral 2.867s) (Fig. 5f). Both the paw ipsilateral to the virus injection and animals infected with a control Cre-dependent mCherry virus showed no change in mechanical/thermal sensitivity, confirming this effect is limited to the targeted hemisphere (Fig. 5d-f). Excitation of this circuit with an excitatory DREADD produced no effect on tactile sensitivity irrespective of the concentration of CNO used, suggesting that it is only the inhibition of this circuit that specifically governs sensitivity to somatic stimulation (Fig. 5d-f and Supp. Fig. 5e).

To confirm that it is the inhibition of the input from axons projecting from S2 into the M2 cortical region that is responsible for the increase in mechanical and heat sensitivity, we inserted cannulas into M2 in inhibitory DREADD animals, and locally microinjected CNO (as used in: ^60,61^) (Fig. 5g, h). Local microinjection of CNO (300nl of 300μM) in M2 increased behavioral mechanical sensitivity similar to that produced by systemic (i.p. 3mg/kg) injections (300µM CNO 0.4778g vs. Saline 0.9243g) (Fig. 5i) and also had an effect on heat sensitivity (300µM CNO vs. 6.031s Saline 10.48s) (Fig. 5j). This indicates that it is the output from S2 to M2 that influences mechanical and thermal nociceptive threshold sensitivity.

The secondary motor cortex is suggested to be involved with the planning of motor behavior in addition to modulating behavioral responses ^44,62^. The increased withdrawal response sensitivity to mechanical and thermal stimulation observed on inhibiting S2 projections to M2 could therefore, be due to an altered motor reactivity. To ascertain whether motor behavior was significantly altered by changing activity of S2-to-M2 neurons, we analyzed gait and motor function of mice infected with DREADDs. Examining the sciatic functional index (SFI) to compare the position of one hindpaw (DREADD affected) to the other (control) during locomotion, we found no significant differences in their locomotory behavior (Supp. Fig. 6a). There was also no change in stride length (Supp. Fig. 6b). This suggests that the S2 to M2 circuit is primarily sensory in nature and exerts its effects on an animals response to sensory stimulation independent of motor planning and learning.

## Discussion

The central neural substrates of somatosensory behavior and how they work together to orchestrate both simple and complex sensory experiences, is still largely unknown. Top-down circuits, often higher order central to lower order central/peripheral circuits, are important modulators of behavior. In somatosensory circuits, top-down modulation can occur from cortical circuits (primary somatosensory cortex (S1), anterior cingulate cortex (ACC)) and subcortical circuits (amygdala and brainstem)^4,63-65^. We previously characterized a S1 excitatory corticospinal circuit in which inhibition produces decreased mechanical sensitivity^4^. However, we noted that this circuit failed to respond to or alter heat-evoked thermosensory behaviors^4^. Indeed, recent work has confirmed this showing that S1 primarily encodes cooling but not heating^6^. We now show that the secondary somatosensory cortex (S2) can fulfill this role and also modulate mechanical responses in a distinct manner from S1. Fiber photometry recordings demonstrate that S2 can respond to mechanical, heat, and cooling applied to the hindpaw. Yet, optogenetic inhibition of S2 produces mechanical and heat hypersensitivity without affecting cooling sensitivity. Other work has identified the posterior insular cortex as a crucial substrate of thermosensory behaviors^6^. While close in anatomical space and sharing some connectivity, whether they interact to achieve a common goal or act on common downstream elements to exert an influence on behavior is an interesting future direction.

Previously defined somatosensory cortical circuits which mitigate sensorimotor action originate from the primary somatosensory cortex (S1)^4,66^ and exert action via direct or indirect spinal connections or through connections with the primary motor cortex (M1)^67^, or from S2 to S1/M1^13,14,36^. However, we now find that the secondary motor cortex (M2) is an important cortical substrate that S2 acts through. Indeed, chemogenetic inhibition of S2-to-M2 neurons phenocopies the effects of broad S2 inhibition. Further, local inhibition of S2 axons that project into M2 also increase mechanical and heat sensitivity. M2 (also known as the supplementary motor area (SMA) or agranular cortex (AGm)) is part of the rodent prefrontal cortex and involved with motor planning, choice-based behavior, and motor learning^44,62,68,69^. Indeed, lesioning of M2 in rodents produces reduced performance in many sensory-based Go/No-Go tasks^41-43,70^. Further, recording neural activity in rats trained in a modified T-maze in which each arm has different probabilities of reward has revealed that M2 displays the earliest choice-related activity of any cortical region^42^. Work in the primate and the mouse whisker system has identified S2 as important for decision making^14,71^. Specifically, S2 to S1 projections are thought to be important in the encoding of choice following a stimulus^14^. It is likely that these three neural substrates work together to process sensory information and produce behavioral responses. However, our behavioral assays are intrinsically different in that the choice (paw withdrawal or not) is not a learned behavior nor coupled to any predictable stimulus. This indicates that S2 inputs to M2 may gate an animal’s behavioral response to defined somatosensory stimuli, in addition to its more complex roles in sensory decision making, choice, association, and learning. Our anatomical tracing supports this theory by demonstrating that layer Va pyramidal neurons in S2 predominantly provide input to layer II/III neurons in M2, a connectivity pattern typically associated with modulating rather than driving cortical responses^52^.

Taken together, this places S2/M2 circuitry as a core mediator of somatosensory behavior. How this circuit works in collaboration with other defined cortical circuits is an interesting future direction, but we hypothesize since distinct areas of the somatosensory cortex appear to process distinct modalities that these circuits operate in a parallel fashion to provide a comprehensive picture of the somatosensory environment.

## Acknowledgements

We would like to thank Mark L. Andermann, Lee B. Barrett, Nick Andrews, Yu-Ting Cheng, Mark Scimone, Jonathan M. Szeber, and David Yarmolinsky, for experimental expertise and feedback. Funding was provided by Charles Robert Broderick III Phytocannabinoid Fellowship Award (D.G.T), William Randolph Hearst Fund Fellowship (Q.J.), NIH R01EY013613 (C.C.), Tan Yang Center for Autism Research (C.C. and Q.J.), R01AT010779 (Z.H.), and BRAIN Initiative Grants funded by NCCIH/NIMH R01 AT011447 (C.J.W) and NINDS R0NS1109947(Z.H.). In addition, we would like to thank the Harvard Neuroimaging Facility (NINDS P30 Core Center Grant #NS072030 and NIH OD 1S10OD026866), the Boston Children’s Hospital Cellular Imaging Core (5P50HD105351-02), the Boston Children’s Hospital Human Neuron Core (IDDRC: U54HD090255), and the Boston Children’s Hospital Viral Core (NEI P30 grant: 5P30EY012196).

## Author Contributions

D.G.T and C.J.W conceived of the experiments, design, and statistical analysis, and wrote the manuscript. D.G.T. and F.P. performed calcium imaging, behavioral experiments, viral tracing, and histology. Q.J. and C.C. performed slice electrophysiology. K.Y., C.C., A.C., performed histology and projection analysis. M.R.B. performed 2-photon serial tomography. M.E. performed RNAscope. J.S. performed surgeries. All authors aided in manuscript editing and writing.

## Figure Legends

**Supplemental Figure 1:**
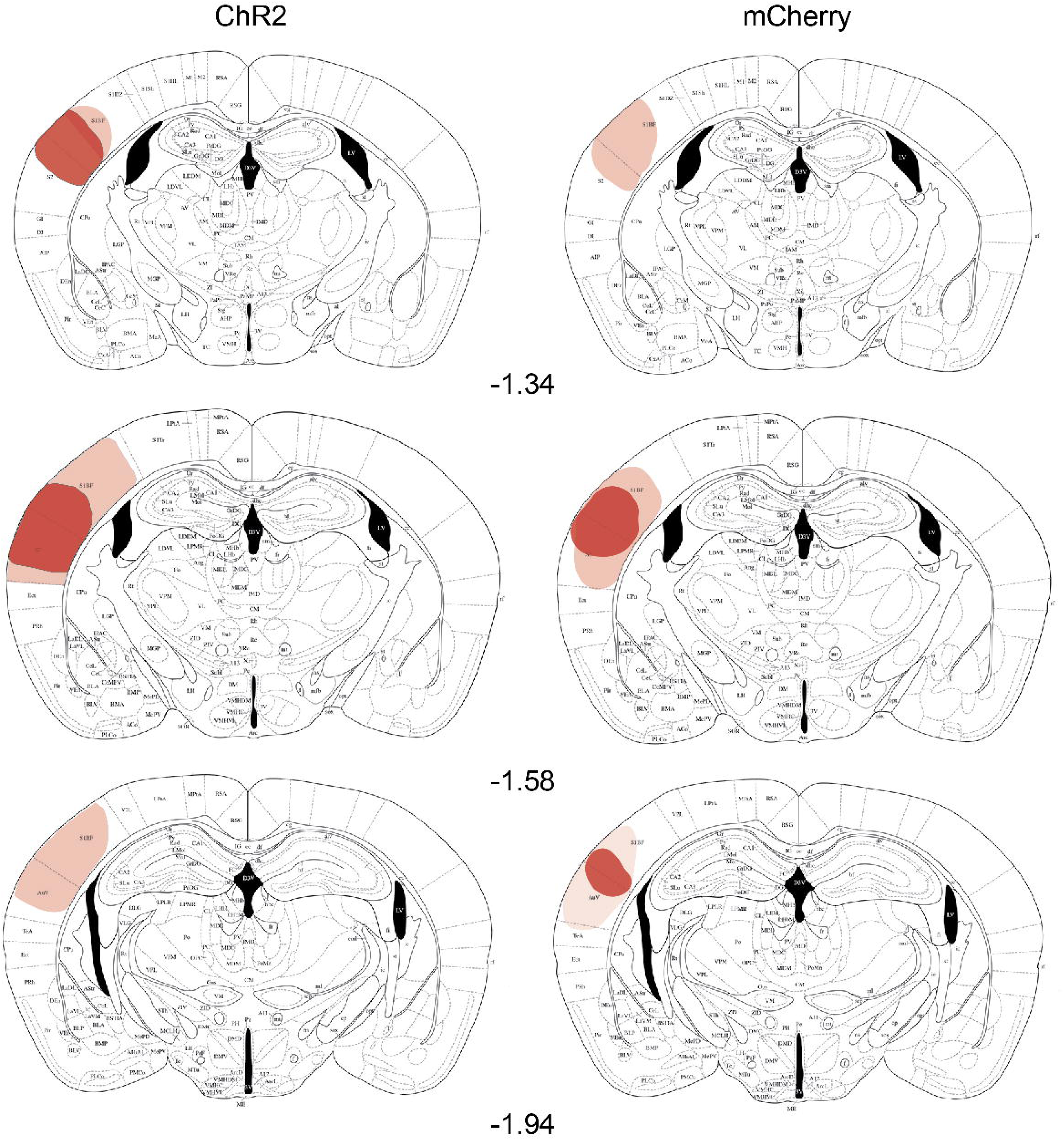
Channelrhodopsin and mCherry viral injection coverage in PV-Cre mice. Averaged spread of virus injection in mice across different anterior-posterior ranges. Darker red indicates more intense expression whereas lighter red is lower expression.

**Supplemental Figure 2:**
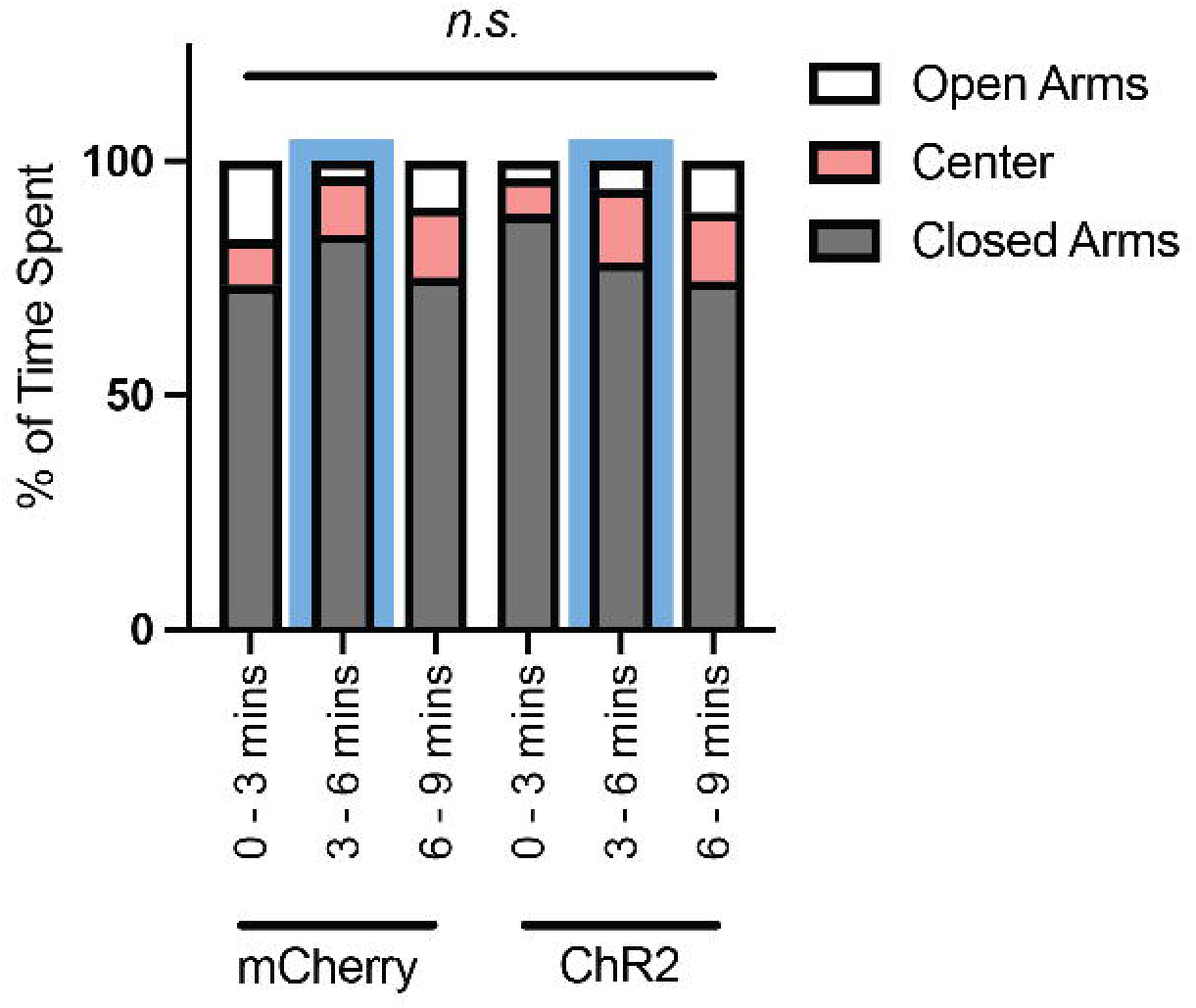
S2 inhibition does not induce anxiolytic behavior. Elevated plus maze total results examining the % of time spent in the closed, center, and open arms. A baseline period was recorded from 0-3 mins, blue light illumination (S2 inhibition) occurred between 3-6 mins, and a recovery period from 6-9 mins. No significant differences observed between mCherry control (n=7) and ChR2 mice (n=6). Two Way ANOVA with Sidak’s.

**Supplemental Figure 3:**
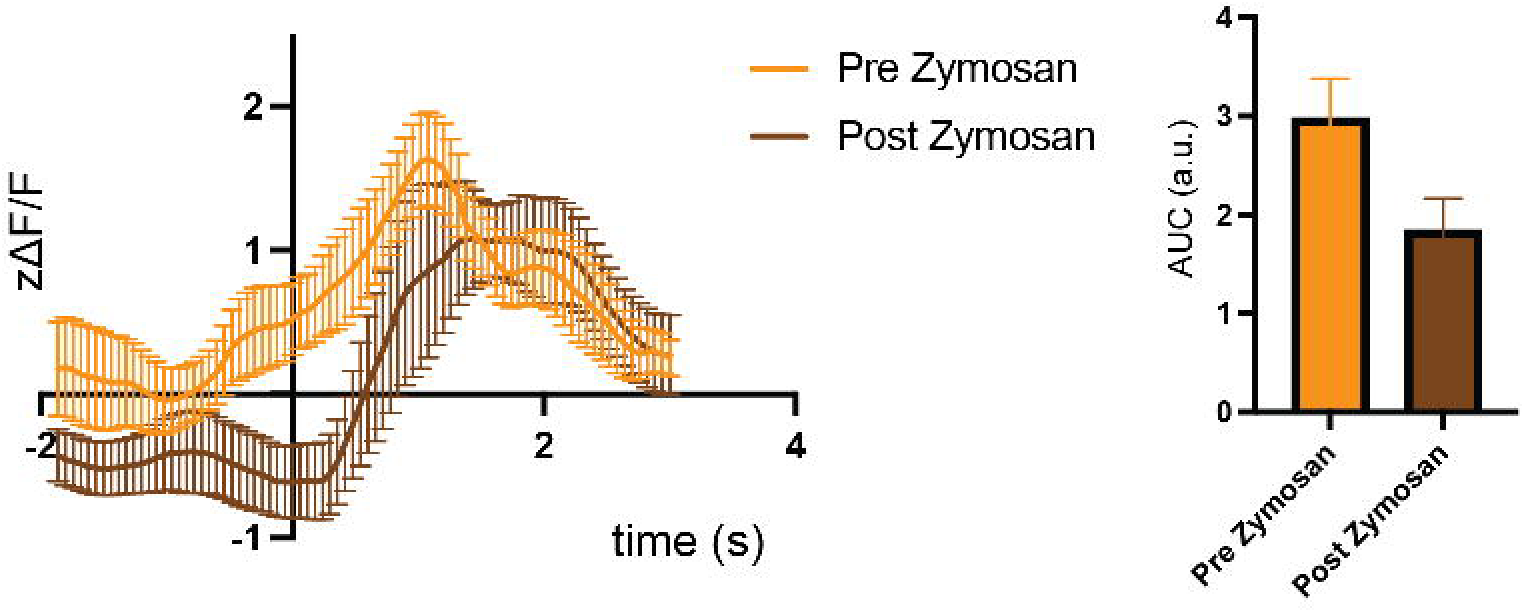
Calcium dynamics of heat responses in parvalbumin interneurons in S2 following hindpaw inflammation. Left: Calcium responses in PV neurons in S2 plotted by Z-scored delta F/F to a heat ramp stimulus at baseline and 4 hours following inflammation induction with zymosan. Time 0 is time of paw withdrawal from the heat source. Right: Area under curve analysis of calcium transients.

**Supplemental Figure 4:**
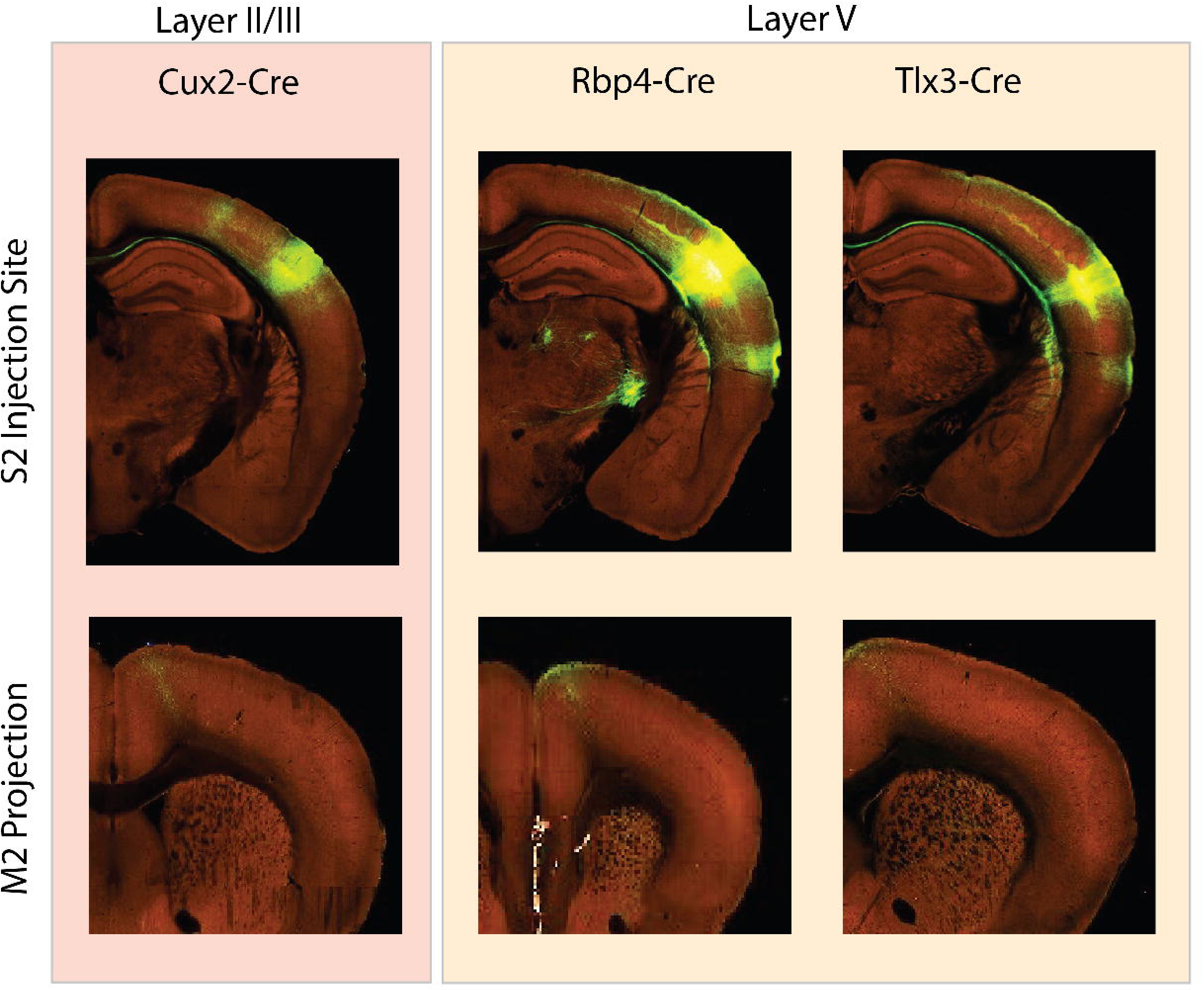
Layer specific contributions of S2 projections to M2. Representative images from the Allen Brain Connectome Atlas (https://connectivity.brain-map.org)38,50. AAV-GFP was injected into S2 of various cre lines (Cux2 for layer II/III and Rbp4 or Tlx3 for layer V) and projections to M2 are clearly identified with largest portion occurring from Rbp4-Cre animals.

**Supplemental Figure 5:**
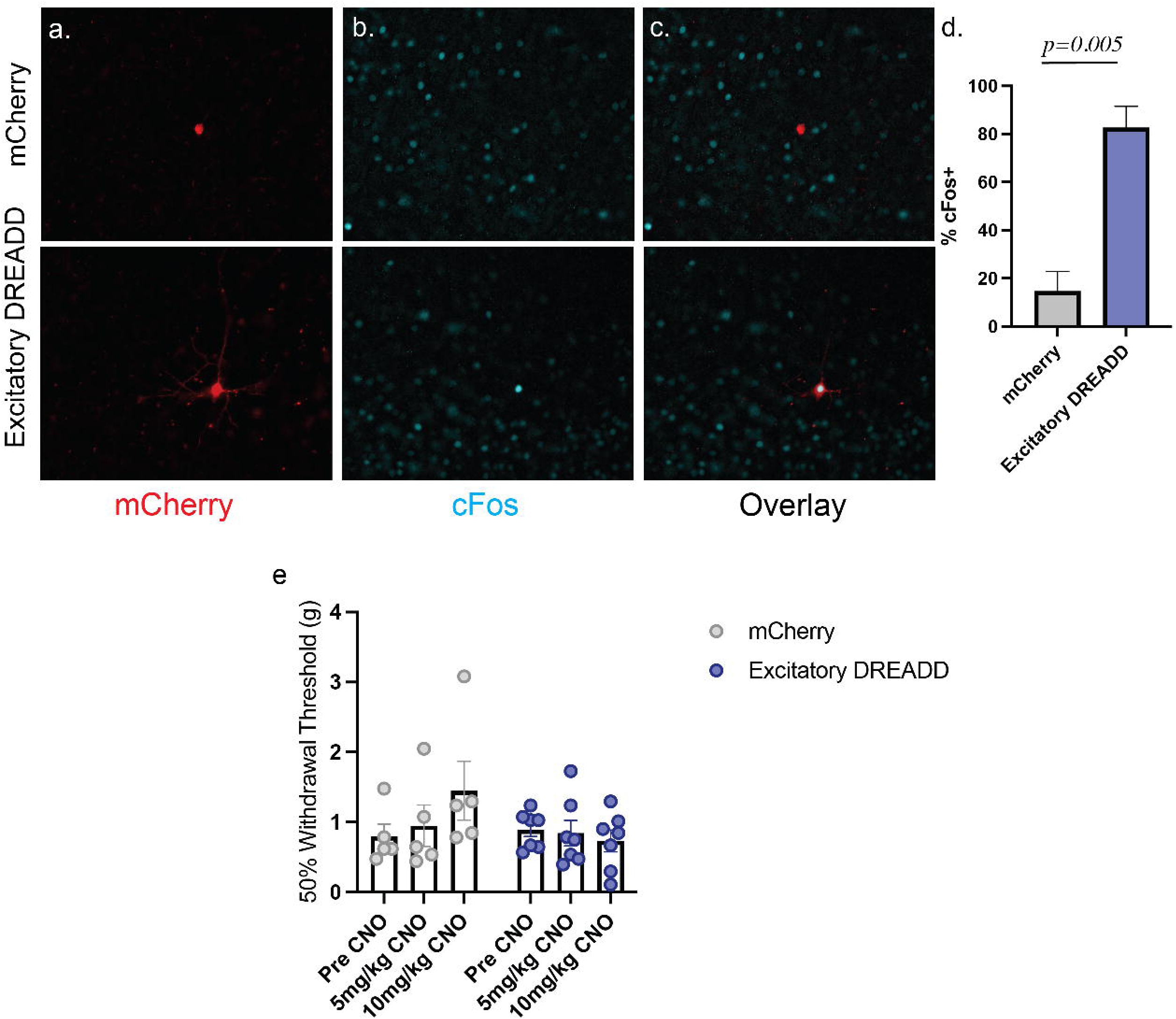
CNO administration increases cFos expression in excitatory DREADD labelled S2-to-M2 neurons but does not influence somatosensory behavior. (A): Representative images showing S2-to-M2 virally labeled neurons with AAV-hSyn-DIO-mCherry or AAV-hSyn-DIO-Hm3q-mCherry (excitatory DREADD). (B): cFos staining in the same plane as in A. (C): Overlay between mCherry labeled neurons and cFos. (D): Quantification of % of mCherry-positive neurons that colabel with cFos. (n=3 animals per group, unpaired t-test). (E): Effect of different doses of clozapine-n-oxide (CNO) on mCherry control and S2-to-M2 excitatory DREADD animals (Two Way ANOVA with Tukey’s, n.s.). Data presented as mean ± SEM.

**Supplemental Figure 6:**
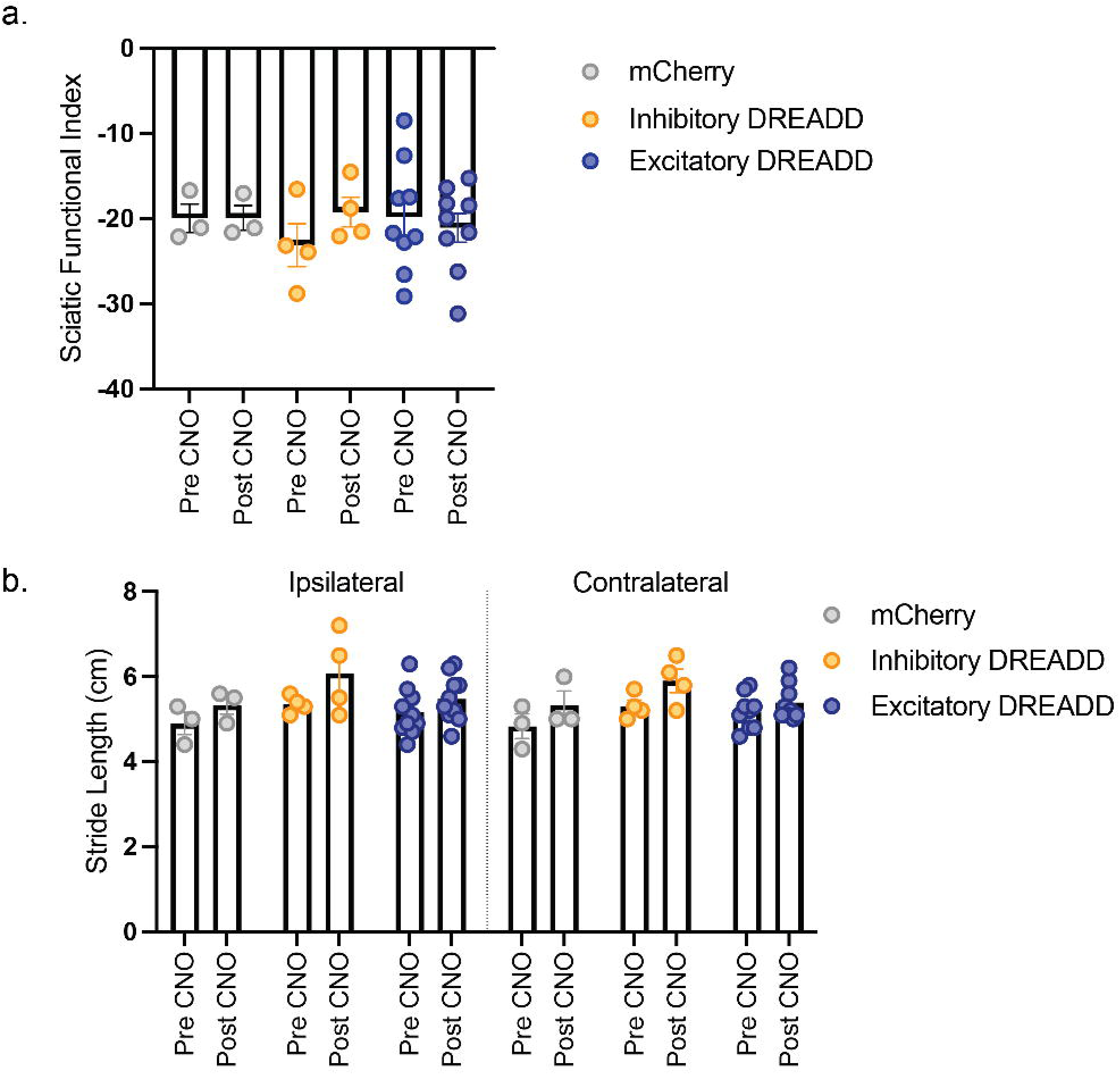
S2 inhibition does not alter motor behavior. (A): Hindpaw placement during locomotion (sciatic functional index) was unaffected in S2-to-M2 chemogenetically inhibited or excited mice. mCherry n=4, Inhibitory DREADD n=5, Excitatory DREADD n=9. Two Way ANOVA. (B): Stride length was unaffected in S2-to-M2 chemogenetically inhibited or excited mice. mCherry n=4, Inhibitory DREADD n=5, Excitatory DREADD n=9. Two Way ANOVA. All data presented as mean ± SEM.

## Methods

### Animals

Both male and female C57BL6/J (Jax #000664) and B6.129P2-*Pvalb^tm1(cre)Arbr^*/J (Jax #017320) mice of roughly equal numbers were used for the behavioral and imaging experiments unless otherwise stated, see figure legends for details. Mice were injected with AAVs between 10-14 weeks of age and behavior was conducted between 3-10 weeks post injection, depending on viral expression. All experiments were conducted in a blinded fashion with strict accordance to the guidelines set forth by the Boston Children’s Hospital Institutional Animal Use and Care Committee under protocol 00001507, 00001546, and 20-05-4165.

### Stereotaxic Injection of Adeno-Associated Viruses and Fiber/Cannula Implantation

All coordinates were identified relative to the bregma on the skull. Coordinates for the hindlimb secondary somatosensory cortex (S2) (ML: 3.9 AP: –1.3 DV: 2.5) were determined based upon our previous anatomical studies ^4^. Secondary motor cortex (M2) coordinates (ML: 0.4 AP: +1.34 DV: 0.8) were chosen based upon the densest projection site from S2 in preliminary anatomical tracing studies. All injection sites were verified post-mortem.

For optogenetic stimulation, 2mm, 200µM diameter 0.39NA optical fiber cannula (Thor Labs CFMLC12L05) and for fiber photometry, 3mm cleaved, 200µM diameter 0.37NA black ceramic fiber optic cannulae (Neurophotometrics Ltd.) were implanted into the S2 region and affixed to the skull with dental cement. Two skull screws (Fine Science Tools 19010-10) were implanted on the opposite hemisphere of the fiber cannulae, roughly above the coordinates of the primary motor cortex and primary visual cortex.

For cannula studies, a 0.8mm cannula was implanted in the M2 region and affixed to the skull with dental cement. Skull screws were used to stabilize the implant as above. A cap that consisted of a 1mm dummy cannula was used when the injector unit was not in use.

For rabies tracing studies, either AAV2/9-Syn-Flex-TVA-oG-GFP or AAV2/8-Syn-Flex-TVA-oG-GFP was injected into the secondary motor cortex of either Penk-Cre (Jax # 025112) or Rbp4-Cre (MGI:4367067) animals at a depth of 200 (to target layer II/III) or 400 micrometers (to target layer V), respectively. Six weeks post-injection, SADdg-EnvA-mCherry was injected at four points along M2 to capture a wide breadth of starter cells. Animals were taken for histology one week following.

Viruses used in this study include: AAV2/9.Syn.Flex.GCaMP6f.WPRE.SV40 (1E+13 Addgene 100833 – 100nl), AAV2/1-CAG-FLEX-rev-ChR2-tdtomato (1.23E+13 gc/mL – Boston Children’s Hospital Viral Core – 125nl), AAV2/retro-CAG-Cre-WPRE (2.89E+13 gc/mL – Boston Children’s Hospital Viral Core – 125nl), AAV2/1-Syn-DIO-hM4Di-mCherry (Boston Children’s Hospital Viral Core – 125nl), AAV2/9-hSyn-DIO-Hm3D(Gq)-mCherry (6.21432E+13 gc/mL – Boston Children’s Hospital Viral Core – 125nl), AAV2/9-hSyn-DIO-mCherry (Boston Children’s Hospital Viral Core) – 125nl, AAV2/9-CAG-tdTomato-WPRE (1.0234E+13 gc/mL Boston Children’s Hospital Viral Core – 50nl), AAV2/retro-hSyn-tdTomato-WPRE (Boston Children’s Hospital Viral Core – 125 nl), SADdg-EnvA-mCherry (2.26E+08 to 4.10E+10 TU/mL Boston Children’s Hospital Viral Core), AAV2/9-Syn-Flex-TVA-oG-GFP (1e13 gc/ml Boston Children’s Hospital Viral Core) AAV2/8-Syn-Flex-TVA-oG-GFP (1e13 gc/ml Boston Children’s Hospital Viral Core).

### Electrophysiology

To obtain brain slices containing S2, mice were anesthetized using isoflurane and decapitated into oxygenated (95% O_2_; 5% CO_2_) ice-cold cutting solution (in mM): 130 K-gluconate, 15 KCl, 0.05 EGTA, 20 HEPES, and 25 glucose (pH 7.4 adjusted with NaOH, 310-315 mOsm). The brain was then removed quickly and immersed in the ice-cold cutting solution for 60 seconds. Coronal slices containing S2 were sectioned and collected. The brain was cut with a steel razor blade, then sectioned into 250 μm-thick slices in the oxygenated ice-cold cutting solution using a sapphire blade (Delaware Diamond Knives, Wilmington, DE) on a vibratome (VT1200S; Leica, Deerfield, IL). The slices collected were allowed to recover at 30°C for 20 minutes in oxygenated saline solution (in mM): 125 NaCl, 26 NaHCO_3_, 1.25 NaH_2_PO_4_, 2.5 KCl, 1.0 MgCl_2_, 2.0 CaCl_2_, and 25 glucose (pH 7.4, 310-315 mOsm).

To detect the inhibitory innervation of PV interneurons on pyramidal neurons in S2 L2/3 or L5, ChR2-mCherry was expressed in PV interneurons and pyramidal neurons were recorded in response to stimulation of PV interneurons. Pyramidal neurons in S2 L2/3 and L5 were visualized through a monitor with projection from the camera of a DIC-equipped microscope (Prime BSI, Teledyne Photometrics). Inhibitory post-synaptic currents (IPSCs) were obtained by holding the membrane potential at 0 mV and with the presence of NBQX (10 μM, 0373, Tocris) and CPP (20 μM, 0247, Tocris) to block AMPA and NMDA receptors, respectively. Evoked IPSCs (eIPSCs) were induced by applying a train of single pulses (0.2 ms) of full-field illumination of blue light through the 60x objective (Olympus LUMplanFL N 60x/1.00W) with interval of 250 ms. The blue light (470 nm, 83 mW/mm^2^) was supplied by a CoolLED pE unit. Spiking of pyramidal neurons were detected through current clamp mode by injecting currents ranging from 100 to 600 pA. The effect of PV inhibitory transmission on the firing of pyramidal neurons was examined by stimulating PV interneurons using blue light at a frequency of 40 Hz with each pulse lasting for 20 ms. Electrophysiological data acquisition and offline analysis were performed using custom software in IgorPro (Wave-Metrics, Portland, OR). eIPSCs were averaged from 3-5 traces.

To detect the expression of inhibitory DREADD, pyramidal neurons in S2 L2/3 or L5 projecting to M2 were recorded through whole-cell patch clamp and tested with CNO. The pyramidal neurons were labeled by mCherry and can be visualized through fluorescence microscope (Olympus, Japan). Glass pipettes (Drummond Scientific) were pulled on Sutter p87 Flaming/Brown micropipette puller (Sutter Instruments) and filled with internal solution containing (in mM): 150 K-gluconate, 8 KCl, 10 EGTA, 10 HEPES (pH7.3, 290-300 mOsm) to optimize the pipette resistance to be 3.5-4.0 MOhm. Patch recordings were performed using a MultiClamp 700B (Axon Instruments, Foster City, CA) and digitized at 20 kHz with an ITC-18 interface (Instrutech). Intrinsic cellular properties of pyramidal neurons were measured in current clamp mode. I-V curves were obtained by recording firing rates when using current injection from 0 to 600 pA in steps of 50 pA with duration of 1 sec. CNO (5 μM, BML-NS105, Enzo) was then perfused into the bath solution. More I-V curves were collected during (20 min since CNO perfusion) and after the washing out of CNO. Four repeats were conducted from four mice.

### Histology

Mice were perfused transcardially with 4% paraformaldehyde (PFA) in PBS. Brains were isolated and fixed overnight in 4% PFA before storage in 1 X PBS. Brains were sectioned with a vibratome between 60-100µm or a cryostat at 30µm and mounted on slides (Fisher Permafrost). Vibratome sections were permeabilized with 1 X PBS with 0.2% Triton-X100, mounted on slides, and coverslipped with mounting media containing DAPI. All injection sites were aligned back to the Allen Brain Atlas, injection sites with substantial off-target infection were excluded. In the cannula experiments, signal from the AAV2/9-CAG-DIO-DREADD(h4Dmi) was amplified using anti-mCherry (Abcam: ab167453) at 1:500 by first blocking with 1 X PBS with 10% goat serum and 0.3% Triton X-100 for one hour at room temperature, incubated with anti-mCherry overnight at 4°C, followed by incubation with goat anti-rabbit 568 (Thermo Scientific: A-110011) for one hour at room temperature, and cover slipped with DAPI mounting media. For identification of excitatory cortical neurons in layer II/III, anti-Foxp1 (Abcam: ab16645 1:500) was used with goat anti-rabbit 488 (Thermo Scientific: A-11008). Slides were visualized and imaged with a Nikon Ti-1200.

### RNAscope

Injections of cholera toxin subunit b conjugated to Alexa Fluor 555 (CTB-555) (Thermo Scientific C34776) was injected at 1X 150-200nl in the secondary motor cortex as described above. Two weeks following injection, animals were processed as above but after incubation with PFA were placed into 30% sucrose in 1 X PBS for 2 days. Brains were rinsed in 1 X PBS and snap frozen in optimal cutting temperature compound (TissueTek 4583) and stored at –80°C until processing. Brains were sectioned with a cryostat at 20 micrometers. RNAscope was performed per manufacturers instructions (ACDBio – RNAscope Multiplex Fluorescent V2 Assay). Probes against Ctip2 (413271-C1), Etv1 (557891-C2), Trib2 (514021-C1), and Satb2 (413261-C2) were used and slides coverslipped with DAPI mounting media. Slides were imaged with Olympus VS120 SlideScanning microscope.

### Fiber Photometry

Mice were habituated to the fiber optic cable for one hour each on two separate days. Photometry signals were acquired by alternative illumination with 470nm (GCaMP) and 410nm light (isobestic control) (Neurophotometrics FP3001). Any mice with calcium responses that occurred during whisker deflection (suggesting fiber was placed in barrel cortex) or sound (suggesting fiber was place in adjacent auditory cortex) were excluded and only animals with responses to hindpaw stimulation were used. Tactile stimuli, heat stimuli, and cold stimuli were presented 10, 5, and 10 times per mouse separated by at least 30 seconds in the case of tactile and 1 minute in the case of thermal. Analysis of Z-scored delta F/F was performed in Matlab as described in ^72^.

### Von Frey Mechanical Sensitivity Assay

Mice were placed in individual square chambers on grated flooring to allow access to the hindpaw from the bottom. Mechanical thresholds were acquired using the Up-Down method in which filaments that exert known forces are applied to the hindpaw in a successive order to ascertain the 50% Withdrawal Threshold. Mice were allowed to habituate to the chamber for 1 hour on two separate days in which 5 stimulations of each of the hindpaws with a 0.6g filament was applied to get the mice accustomed to foot stimulation. On the day of sensory testing, mice were allowed to habituate in the chambers for 1 hour before data acquisition.

### Hargreave’s Thermal Assay

Heat sensitivity was assessed by placing mice in individual square chambers on a glass surface heated to 30°C. A radiant heat source targeted to the hindpaw was applied until a withdrawal response or a 30-second cut off was reached. Mice were allowed to habituate to the chamber for 1 hour on two separate days and 1 hour before on the day of testing.

### Acetone Cold Sensitivity Assay

Cold sensitivity was determined by placing individuals in square chambers on grated flooring in order to access the hindpaw from below. Mice were allowed to habituate to the chamber for 1 hour on two separate days. A 30µL drop of acetone was placed on the surface of either the ipsilateral or contralateral paw and the amount of time the mouse spent biting or flinching the paw was quantified.

### Optogenetic Activation of Inhibitory Parvalbumin Neurons

Mice were habituated on two separate days for one hour each with a rotary fiber optic cable connected (Thor Labs RJPFL2) via a ceramic adapter to the optic fiber implant. Blue laser light pulses (470nm, 40 Hz, 3mW output) were administered during the duration of the experiment to activate PV neurons with a maximum exposure of 1 minute per test.

### Thermal Conditioned Place Preference Assay

Two thermal plates (Bioseb BIO-CHP) forming two separate arenas (165mm x 165mm) were enclosed in an acrylic box with one side colored black and the other black-and-white striped. A black door allowed separation between the two arenas during training. Mouse tracking was enabled by an overhead camera that tracked the center point of the mouse and % of time spent in each arena was tracked and calculated (Ethovision). Each plate was set to 39°C. Three weeks post-surgery, mCherry controls and ChR2 injected mice were assigned randomly to receive blue light stimulation (3mW, 40Hz, 1 minute on/1 minute off for 30 minutes) in either the black or white striped chamber. Day 1, mice were habituated to the entire apparatus for one hour. Day 2, mice were pre-tested to determine baseline preference for 15 minutes. Any mouse with a pre-preference of >80% was excluded. Day 3 – 6, mice were trained to associate the stimulus with the chamber by restricting to one chamber and received stimulation in the paired chamber or did not in the unpaired chamber for 30 minutes. On Day 7, place preference was determined by the mouse’s % of time spent in either chamber in a 15-minute time window.

### Elevated Plus Maze

An elevated maze consisting of two open arms, two closed arms, and a center arena was used to assess anxiolytic behaviors as in ^73^. Briefly, mice were placed in the center arena and recorded using Ethovision XT software (Noldus) for 9 minutes. The first three minutes were the baseline period with no light, 3 – 6 minutes consisted of blue light stimulation at the same parameters described above, and 6 – 9 minutes the light stimulation was stopped to assess any after effects. Percentage of time spent in each arm was calculated and statistically compared.

### Analysis of motor behaviors

Motor behaviors in chemogenetic manipulated mice were recorded via Digigait. In brief, mice were placed on an illuminated treadmill with a camera below to capture paw placement. The treadmill was set to 20cm and mice allowed to walk for a 5 minute period. Sciatic functional index and stride length were calculated with the Digitgait Noldus software.

### Two Photon and One Photon Serial Tomography Mapping of S2 Projection Targets

Two mice with 50nl injections of AAV2/8-CAG-tdTomato in S2 were used for serial two photon tomography mapping. Mice were perfused transcardically with 4% paraformaldehyde (PFA), brains isolated and incubated in 4% PFA overnight at 4°C. Brains were then embedded in 4.5% oxidized agarose and coronal sections of 1.38um^2 resolution at 50um optical section were taken using the TissueCyte (Tissue Vision Inc.). Images were aligned back to the Allen Brain Atlas using Neuroinfo software (MBF Bioscience) and projections were manually annotated and quantified using ImageJ. One mouse with a 100nl injection of AAV2/8-CAG-tdTomato in S2 was processed traditionally with a vibratome and imaged with an IXM Confocal microscope to confirm the two photon findings.

### Clozapine-N-Oxide (CNO) administration during behavioral tests

To activate DREADD chemogenetic receptors in behavioral tests, 3 mg/kg clozapine-n-oxide (CNO – Enzo Biosciences BML-NS105) (first dissolved in DMSO and saline added until 0.02% DMSO final solution) was injected intraperitoneally 30 minutes prior to behavioral measurement. All behavioral measurements were separated by at least 24 hours to ensure metabolism of CNO.

### Clozapine-N-Oxide (CNO) local administration during behavioral tests

For local administration of CNO in M2 via cannula, a 33 gauge injector cannula was attached to the cannula pedestal (P1 Technologies C315GS). Mice were habituated to the injector cannula for 2 days prior to behavioral assessment for 30 minutes each. On the day of testing, mice were habituated to the cannula injector for 5 minutes, 300 nanoliters of saline or different concentrations of CNO injected at 100nl/min, and diffusion allowed to proceed for 2 minutes before behavioral assessment

### Validation of Excitatory DREADD via cFos Staining

Three mice with S2-to-M2 neurons labeled with either mCherry or Excitatory DREADD were injected with 3mg/kg CNO. Animals were placed in the dark for one hour before perfusion. Histology proceeded as detailed above.

### Zymosan Injection into the Hindpaw for Peripheral Inflammation

Zymosan (20μl 5mg/mL Sigma: Z4250) was injected into the hindpaw and GCaMP fiber photometry was performed as above during von Frey and Hargreave’s assays at baseline and the peak of sensitivity (4 hours post injection).

*Statistics:* Statistics were performed in GraphPad Prism version 9. Exact tests performed are listed in the figure legends. For the entire manuscript, significance is defined as *p=<0.05, **p=<0.005, ***p=<0.0005 unless otherwise stated.

